# Bee pasture as a buffer against flower-mediated disease transmission

**DOI:** 10.1101/2024.04.24.590937

**Authors:** Jomar Fajardo Rabajante

**Affiliations:** Institute of Mathematical Sciences and Physics, University of the Philippines Los Baños, 4031, Laguna, Philippines

**Keywords:** flower sharing, niche overlap, optimal foraging, wildflower strip, pathogen spillover

## Abstract

Bees play a crucial role as pollinators in ecosystems, yet they face the risk of flower-mediated diseases that can spread among their communities. Infected bees can inadvertently transmit infective agents, such as microbial spores, to other bee colonies through shared floral resources during foraging. To address this challenge, we propose a solution involving the strategic planting of bee pasture as buffer zones around foraging areas, coupled with the careful placement of beehives to segregate bee colonies. However, this strategy presents dual potential outcomes: the buffer zone could act as a protective barrier, reducing disease transmission; or attract more bees from other colonies, thereby increasing infection risk. Employing mathematical modeling, we explore the intricate dynamics of this strategy to minimize negative outcomes, considering factors such as colony strength, optimal foraging behavior, and foraging area locations. Our results recommend implementing the following measures to safeguard the target bee colony against disease: (i) ensuring ample food sources are available in the vicinity of the target bee colony, and (ii) establishing bee pasture in a distinct, outlying area to serve as a buffer zone. While the bee pasture buffer zone cannot guarantee complete immunity from infection, it can effectively delay the spread of inter-colony diseases. This proposed strategy should be combined with other sound beekeeping practices for comprehensive disease management. Our analysis aims to provide insights into optimizing practices to protect the health of both managed and wild bee populations, and bolster ecological resilience.

**Significance Statement:** Bees, vital pollinators in ecosystems, face the risk of flower-mediated diseases transmitted between colonies. Flower-mediated disease transmission occurs when bees spread pathogens to each other through sharing of flowers during foraging. To mitigate this, we propose planting bee pasture buffer zones around foraging areas. Using mathematical modeling, we recommend ensuring nearby food sources for the target bee colony and establishing bee pasture buffers. While not guaranteeing immunity, this strategy can delay disease spread. Combining this with other beekeeping practices can comprehensively manage diseases. This research underscores the importance of bee pasture areas for protecting both managed and wild bee populations and strengthening ecosystem resilience.

**MSC Classification:** 92-10, 92D40, 92D30

## 1. Introduction

Pollinators, such as bees, play an important role in maintaining the ecological balance of various ecosystems by facilitating the reproduction of flowering plants. The ecological services rendered by bees are indispensable for biodiversity maintenance and serve as a vital factor in providing food source for numerous animal species [1–4]. In agriculture, bees provide invaluable pollination services, essential for the cultivation of various crops, including high-value ones like apple, coconut, coffee, cucurbits, mangoes, sunflower, among others [5, 6]. However, their pivotal role faces numerous challenges, including the impacts of climate change, colony collapse disorder, pesticide use, and the prevalence of diseases [7–9].

The location of beehives is a critical factor in beekeeping practices [10, 11]. Beehives are typically placed in areas with abundant floral resources to support the foraging activities of bee colonies [10, 12]. Strategic placement of beehives is crucial for maximizing the benefits bees provide, such as effective pollination, while simultaneously mitigating challenges like competition, food scarcity, predation, parasitism, exposure to toxic substances like pesticides, and adverse human practices that are not conducive to bee well-being [1, 7–10, 13].

Bee diseases encompass a variety of pathogens, including bacterial, fungal, and viral agents, which can severely impact health and productivity of bee colonies. These diseases, such as American Foulbrood (bacteria), European Foulbrood (bacteria), Nosemosis (fungus), Chalkbrood (fungus), and Sacbrood (virus) [8, 14–16] can spread through various means, including robbing of infected food sources, drifting from infected hives, swarming, and sharing of beekeeping equipment [15, 17], facilitating the disease’s dissemination to new areas. Additionally, macroparasites like Varroa mites further compound the challenges faced by bee populations [8, 15].

Pathogen spores and other infectious agents may be deposited by visiting pollinators onto flowers within just a few hours of foraging [18–22]. These spores have the potential to remain viable for a long time, and an infected bee can shed significant volumes of them [17, 23]. Hence, one pathway for bee disease transmission is through flower sharing, which is possible through niche overlaps of visiting pollinators. The interaction between infected flowers and foraging bees creates opportunities for disease transmission within and between bee colonies, where visited flowers serve as hubs for the spread of diseases [18, 21, 24]. Understanding the spatio-temporal dynamics of disease transmission mediated by floral resources is essential for developing effective strategies to mitigate its impact on bee populations [25]. The proximity of beehives to each other and to potential sources of infection can influence the transmission dynamics of bee diseases [17]. In response to this challenge, strategic interventions are required to minimize the transmission of these diseases while ensuring the sustainability of bee populations.

In this context, the central question arises: In the presence of forage-related diseases, how can we minimize or delay disease transmission among bee colonies? One potential solution that we propose is the implementation of bee pasture buffer zones around foraging areas, coupled with strategic hive placement to segregate bee colonies. However, this strategy presents dual opposite outcomes: (i) it can serve as a protective barrier, mitigating disease transmission; or (ii) it may inadvertently attract more bees, thereby increasing the likelihood of infection spread.

The success of implementing bee pasture buffer zones hinges on their capacity to yield more favorable outcomes than adverse ones. This emphasizes the need for a straightforward yet comprehensive investigation of the underlying dynamics and the development of refined beekeeping strategies to protect bee health and enhance ecological resilience. To address this, we will utilize mathematical modeling to delve into these dynamics. This research could serve as a support in emphasizing the importance of planting or preserving bee pasture areas [12, 26–31].

## 2 Methods

In the mathematical model, we consider two colonies: Colony A and Colony C (see Figure 1). Colony C serves as the primary source of the disease (akin to a reservoir host population), while Colony A represents the target colony we aim to protect from the disease (potential novel host population). The foraging landscape comprises three distinct areas: *F*_*A*_, the foraging area near Colony A; *F*_*C*_, the foraging area near Colony C; and the buffer zone located within *F*_*B*_. Foragers from Colony A prioritize flower visitation based on distance, with a preference for *F*_*A*_, followed by *F*_*B*_, and finally *F*_*C*_. Conversely, foragers from Colony C prioritize flower visitation in *F*_*C*_, followed by *F*_*B*_, and lastly *F*_*A*_. The spread of disease from Colony C to Colony A occurs via flower-mediated transmission.

**Fig. 1.**
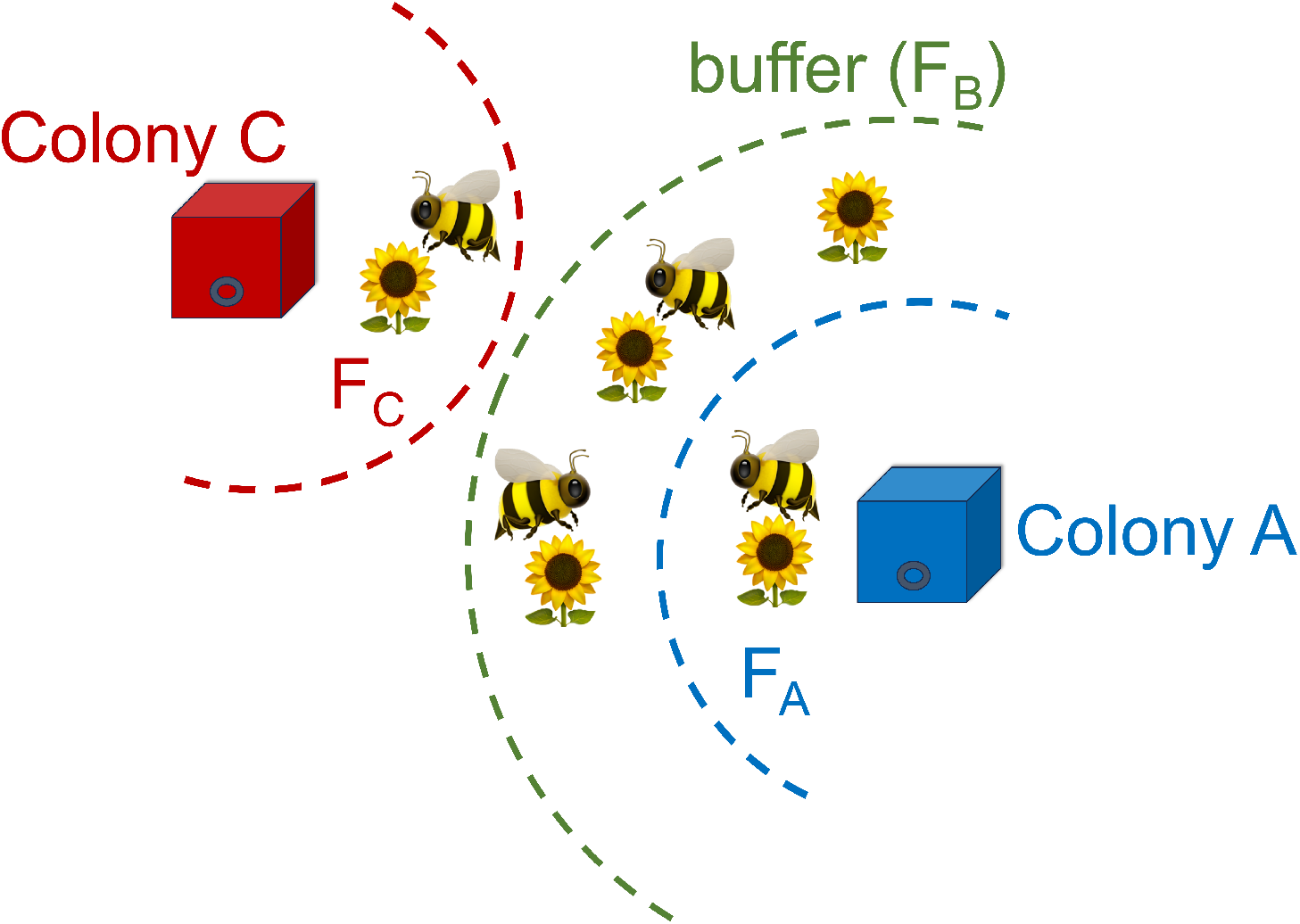
Simplified landscape considered in the mathematical model. There are two (2) colonies: A and C. Colony A is the target colony to be protected, while Colony C is infected. *F*_*A*_ is the foraging area near Colony A, while *F*_*C*_ is near Colony C. *F*_*B*_ is the foraging area (e.g., wildflower strip bee pasture) serving as buffer.

Without losing generality, our model operates under several important simplifying assumptions. First, we assume similar bee species for Colony A and C. Second, the model adheres to the principles of optimal foraging theory, where bees aim to maximize benefits while minimizing costs [32]. Benefit is associated with food source consumption, represented by flower visitation, while cost is associated with flight distance [10, 11, 25]. Third, our model focuses exclusively on inter-colony disease spread, specifically through food source visitation, with Colony C serving as the sole source of the disease. We do not account for intra-colony disease spread within Colony A. Other potential sources of disease transmission, such as sharing of beekeeper equipment, are not included. Fourth, the possibility of dilution effect is taken into account, where the concentration of infectious agents in flowers decreases with a higher density of flowers [33, 34]. Fifth, our model does not consider complex inter-colony food competition between Colony A and C. Sixth, we assume that the number of flowers in each foraging area is dense enough to attract bees [25, 35–38, 38, 39]. Lastly, the model does not incorporate various other constraints that may influence disease transmission dynamics, such as the presence of bee chemical cues, complex barriers, differential floral preferences, floral fidelity, differential length of visitation, differential pathogen load, possible attraction to diseased plants, or the use of optic flow information as an odometer to estimate distance [20, 40–48]. These simplifications are made to streamline the analysis and focus on the core dynamics of disease transmission via food source visitation between colonies A and C, hence, enabling us to derive straightforward yet generalizable conclusions from our model.

For the simulations, we employ the equations provided in the Supplementary Information, derived from the diagram depicted in Figure 2. Let *S*_*A*_ represent the population size of susceptible bees in Colony A, *I*_*A*_ denote the population size of infected bees in Colony A, and *I*_*C*_ signify the population size of infected bees in Colony C. Additionally, let *F*_*i,S*_ (*i* = *A, B, C*) denote the number of flowers in foraging area *F*_*i*_ without infectious agents, and *F*_*i,I*_ represent the number of flowers in foraging area *F*_*i*_ harboring infectious agents.

**Fig. 2.**
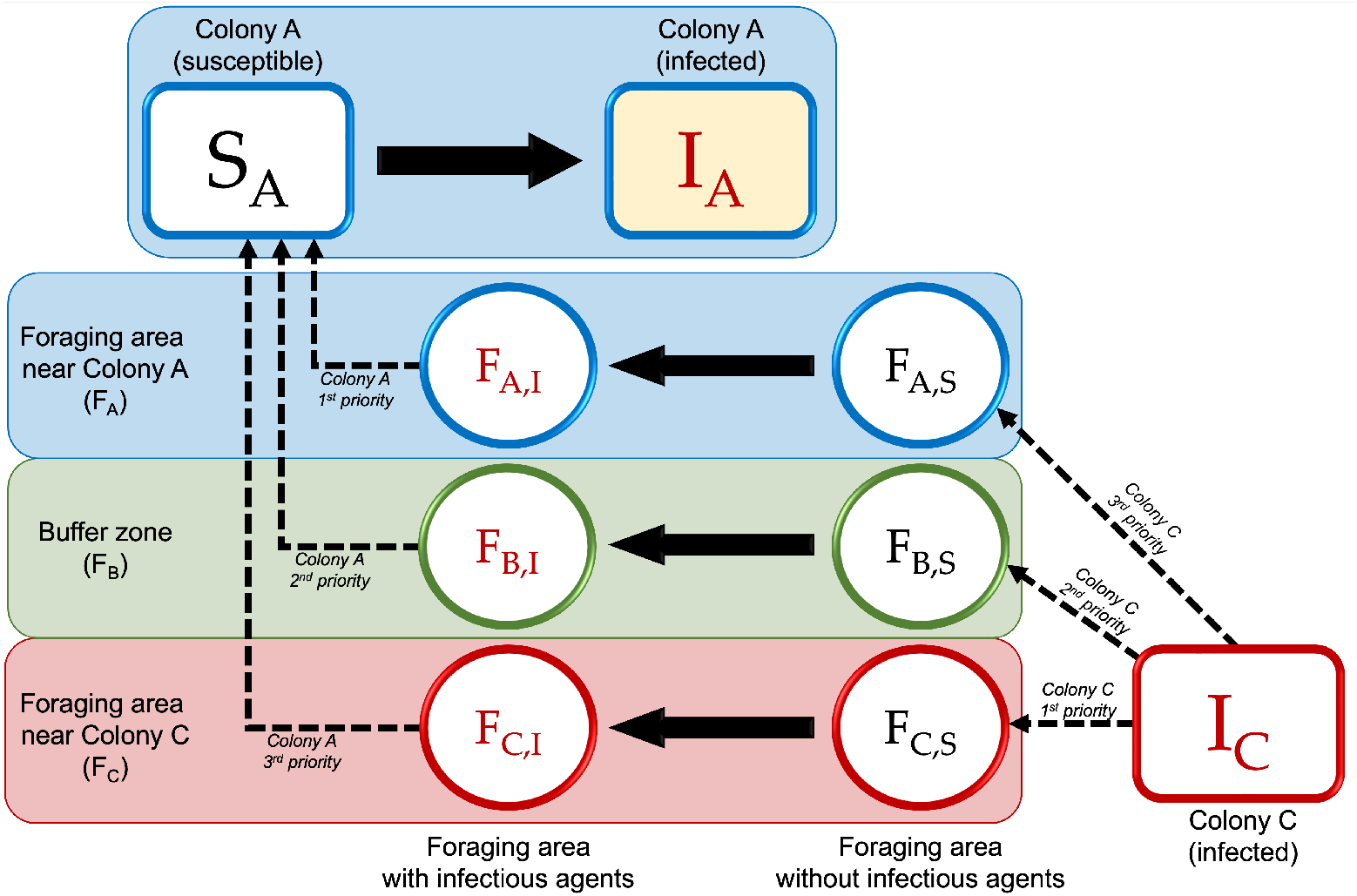
Compartment diagram illustrating the transition of bees within Colony A from a susceptible state to an infected state through the process of visiting flowers contaminated with infectious agents. These infectious agents are initially deposited on the flowers by bees from Colony C and subsequently transmitted to bees in Colony A during their foraging activities.

Foragers from Colony C visit plants in *F*_*A*_, *F*_*B*_, and *F*_*C*_, depositing infectious agents onto the visited flowers. Flowers initially counted in *F*_*i,S*_ that are visited by a bee from Colony C will transition to a new status and will be counted in *F*_*i,I*_, signifying that the flower has been contaminated with infectious agents. Susceptible bees from Colony A (*S*_*A*_) that visit flowers counted in *F*_*i,I*_ will transition to the infected state (*I*_*A*_). However, not all flowers will be visited by bees from both colonies. If there are sufficient food sources in their respective nearby areas to sustain the foragers, they may not need to venture to flowers in distant locations. Nonetheless, as bees from both Colony A and Colony C engage in foraging activities and visit flowers within overlapping foraging areas, the risk of Colony A becoming infected increases. The interaction between bees from different colonies creates opportunities for the spread of diseases, particularly when food sources in surrounding areas are limited.

## 3 Results

It is possible that disease will spread once there is an infected colony and active sharing of food occurs. Therefore, we cannot guarantee zero infection. However, the timing of disease spread may vary. Some cases may exhibit faster spread while others may experience slower transmission. Here, we examine the number of bees in Colony A infected through flower-mediated transmission at *t* = 400. This simulation period spans 400 minutes, representing, for instance, a 6-hour morning foraging period when bees typically forage as inflorescence are in bloom. The lower the number of infected bees in Colony A at *t* = 400, the slower the disease spreads, which aligns with our objective.

In this paper, the size of the foraging area denotes not just the spatial scale but also the density of flowers in the area. It is assumed that each foraging area is distinguishable from each other (e.g., relative to the distance from the colony), and the number of flowers is dense enough to attract bees. In our simulations, we initially demonstrate that the bee pasture buffer zone can yield dual opposite outcomes. As depicted in Figure 3, increasing the number of flowers in *F*_*B*_ (referred to here as ‘buffer zone size’) may initially attract more bees, thereby heightening the risk of infection spread up to a critical threshold (around 10k buffer zone size in the example shown in Figure 3). Beyond this threshold, further increases in the buffer zone size can transform it into a protective barrier, delaying the disease transmission.

**Fig. 3.**
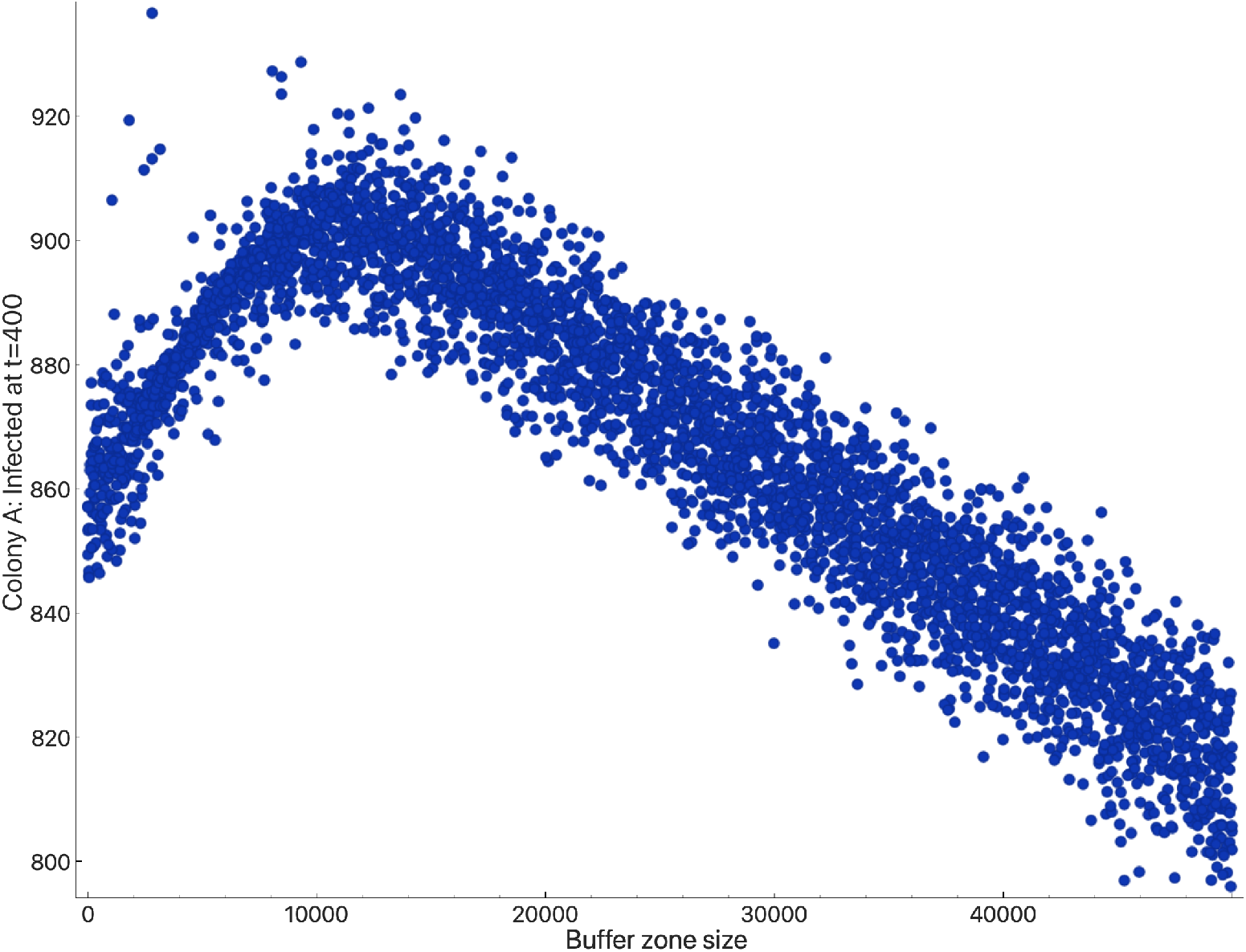
An example showing the dual opposite outcomes of the bee pasture buffer zone. Each point represents the number of bees in Colony A infected through flower-mediated transmission at *t* = 400, given a specified number of flowers in *F*_*B*_ (or ‘buffer zone size’). These points are generated from 4,000 simulation runs.

Next, we assess the strength of Colony C relative to Colony A. Stronger colonies, characterized by a higher number of foragers, are more likely to contribute to flower-mediated disease spread (Figure 4). In some instances, when Colony C is stronger than Colony A, flower-mediated disease transmission can occur more rapidly. Typically, we lack information regarding the characteristics of colonies outside our bee farm (e.g., Colony C). Therefore, it is prudent to prepare for the scenario where Colony C is strong.

**Fig. 4.**
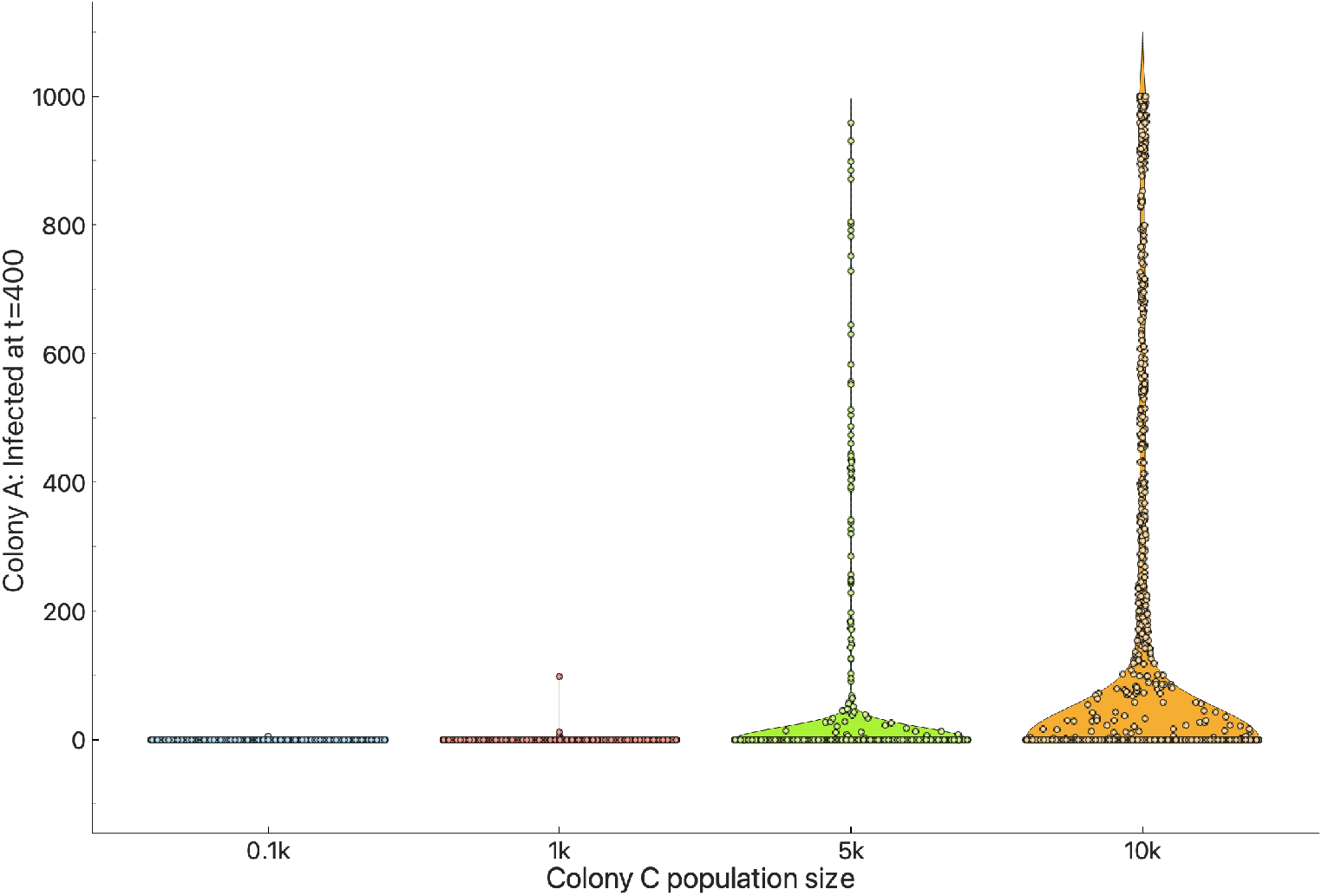
Weak vs Strong Infected Colony C. In this example, Colony A strength is set to a population size of 1k. A stronger Colony C may lead to faster disease transmission. The violin plots display points generated from 8,000 simulation runs (2,000 simulations per plot).

In order to specifically address the question of whether a bee pasture buffer zone can minimize or, at least, delay flower-mediated disease transmission among bee colonies, we need to analyze this proposed strategy by examining the outcomes when we change the number of flowers in the foraging area near Colony A (*F*_*A*_) and in the buffer zone (*F*_*B*_). We specifically focus on these two areas because, in practice, they are the regions that can be controlled by the beekeeper. Typically, we have limited control over the foraging area outside our bee farm.

Figure 5 illustrates the impact of the change in the number of flowers near Colony A (*F*_*A*_). Increasing the number of plant sources in *F*_*A*_ can offer benefits to Colony A in several ways, such as ensuring an adequate food supply to Colony A and enabling beekeepers to confine foragers of Colony A within *F*_*A*_. However, this may not always be advantageous if Colony C has a limited food source in *F*_*C*_ or in *F*_*B*_. In such cases, foragers from Colony C might fly to *F*_*A*_ and inadvertently deposit infectious agents, which is not our desired outcome.

**Fig. 5.**
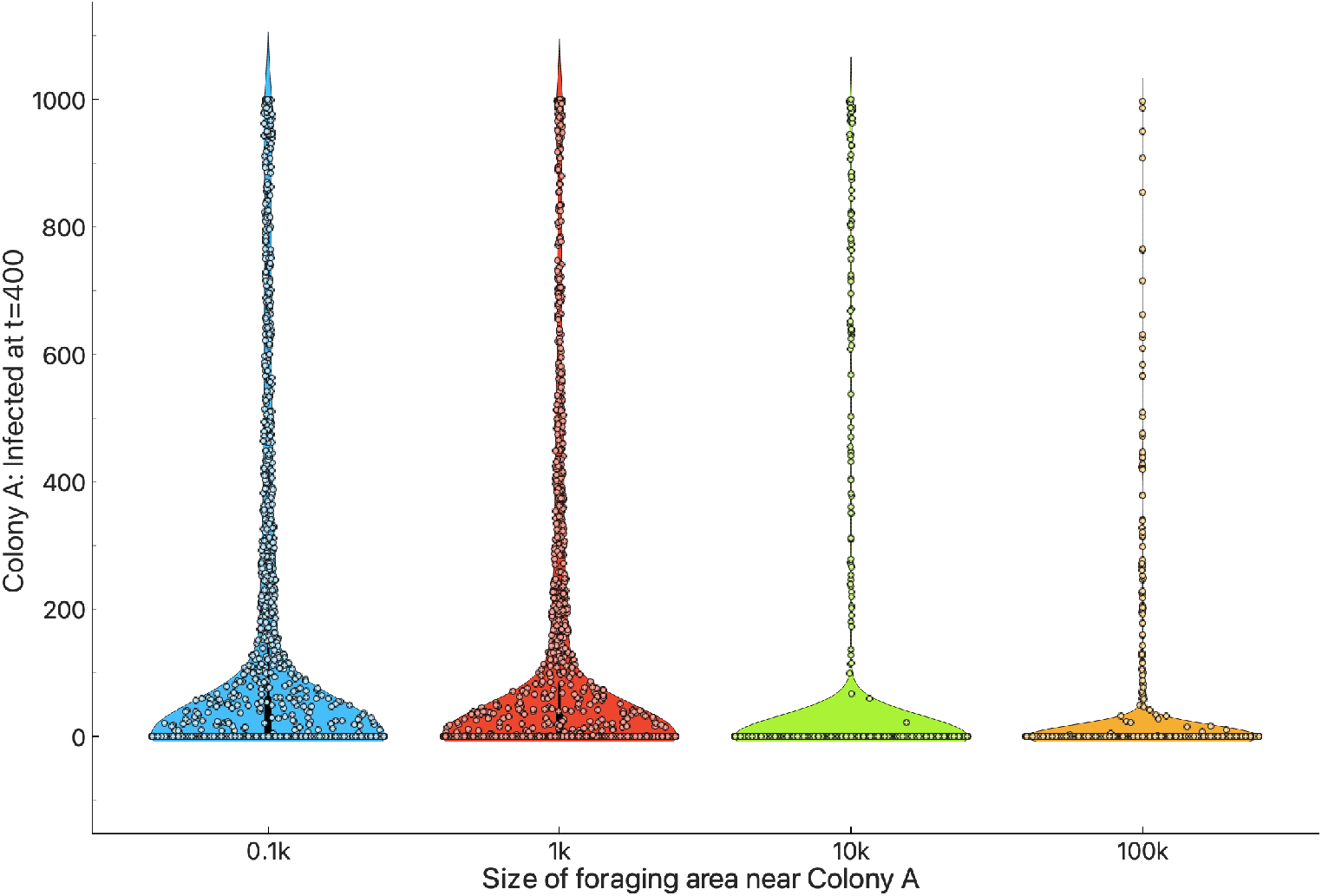
Increasing the number of plant sources in *F*_*A*_ (size of the foraging area near Colony A) does not guarantee slower flower-mediated disease transmission. The violin plots display points generated from 8,000 simulation runs (2,000 simulations per plot).

Figure 6 illustrates the impact of the change in the number of flowers in the buffer zone (*F*_*B*_). While it is not guaranteed to completely prevent flower-mediated transmission, increasing the bee pasture in *F*_*B*_ helps to delay the spread of disease. Providing an ample food source in *F*_*B*_ for Colonies A and C, along with the dilution effect, can mitigate the negative outcome of having a buffer zone.

**Fig. 6.**
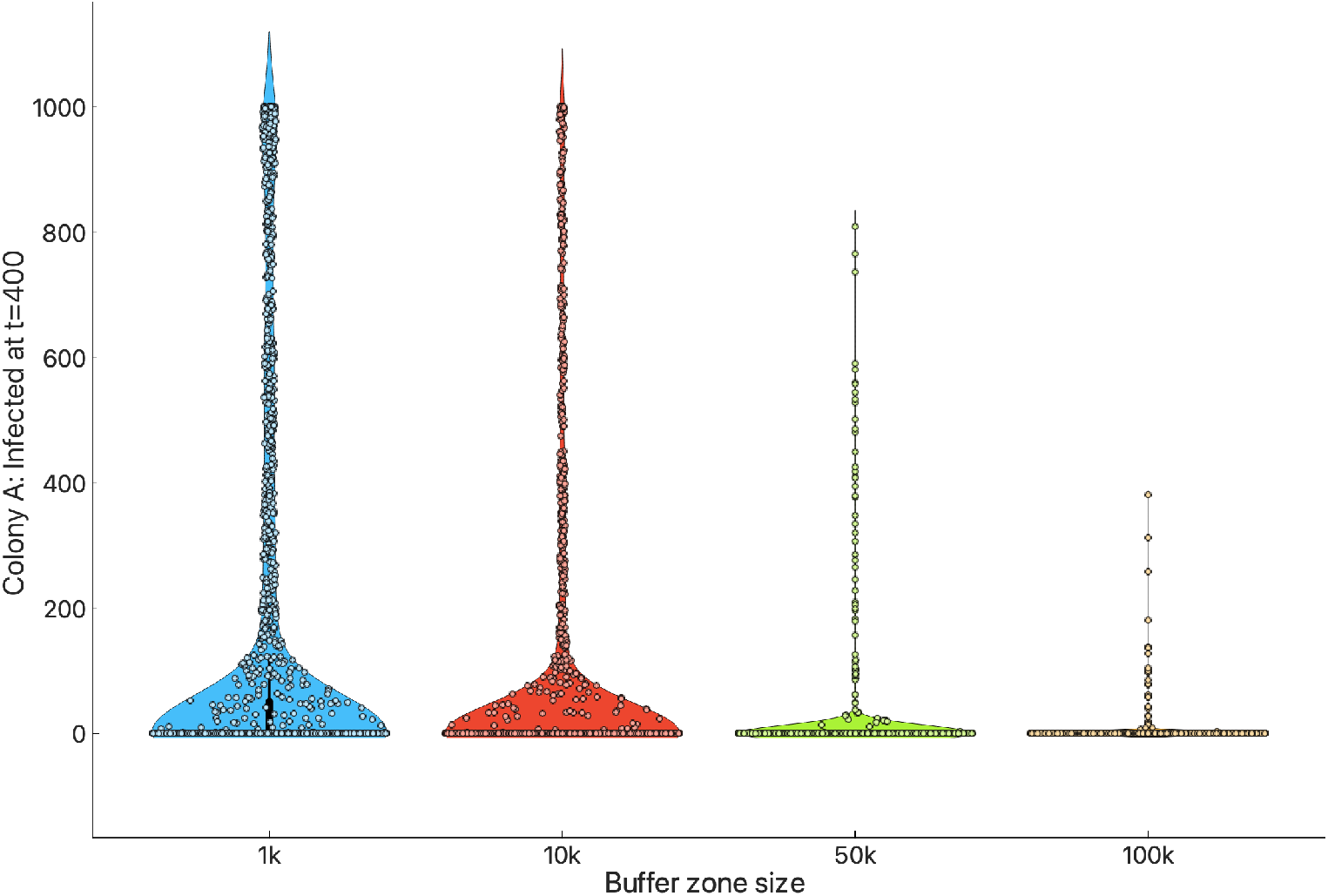
Increasing the number of bee pasture plants in *F*_*B*_ (buffer zone size) contributes to delaying flower-mediated disease transmission. The violin plots display points generated from 8,000 simulation runs (2,000 simulations per plot).

To further investigate the dynamics and elucidate the impact of varying the number of flowers in *F*_*A*_ and *F*_*B*_, we examine Figures 7 and 8. As depicted in Figure 7, it is crucial for Colony A to have an adequate food supply in its nearby foraging area *F*_*A*_; however, this alone may not achieve the desired outcome. It should be complemented with increased bee pasture in the buffer zone. Similarly, the significance of the buffer zone’s impact is evident in Figure 8. Indeed, having sufficient bee pasture in the buffer zone can slow down flower-mediated disease transmission.

**Fig. 7.**
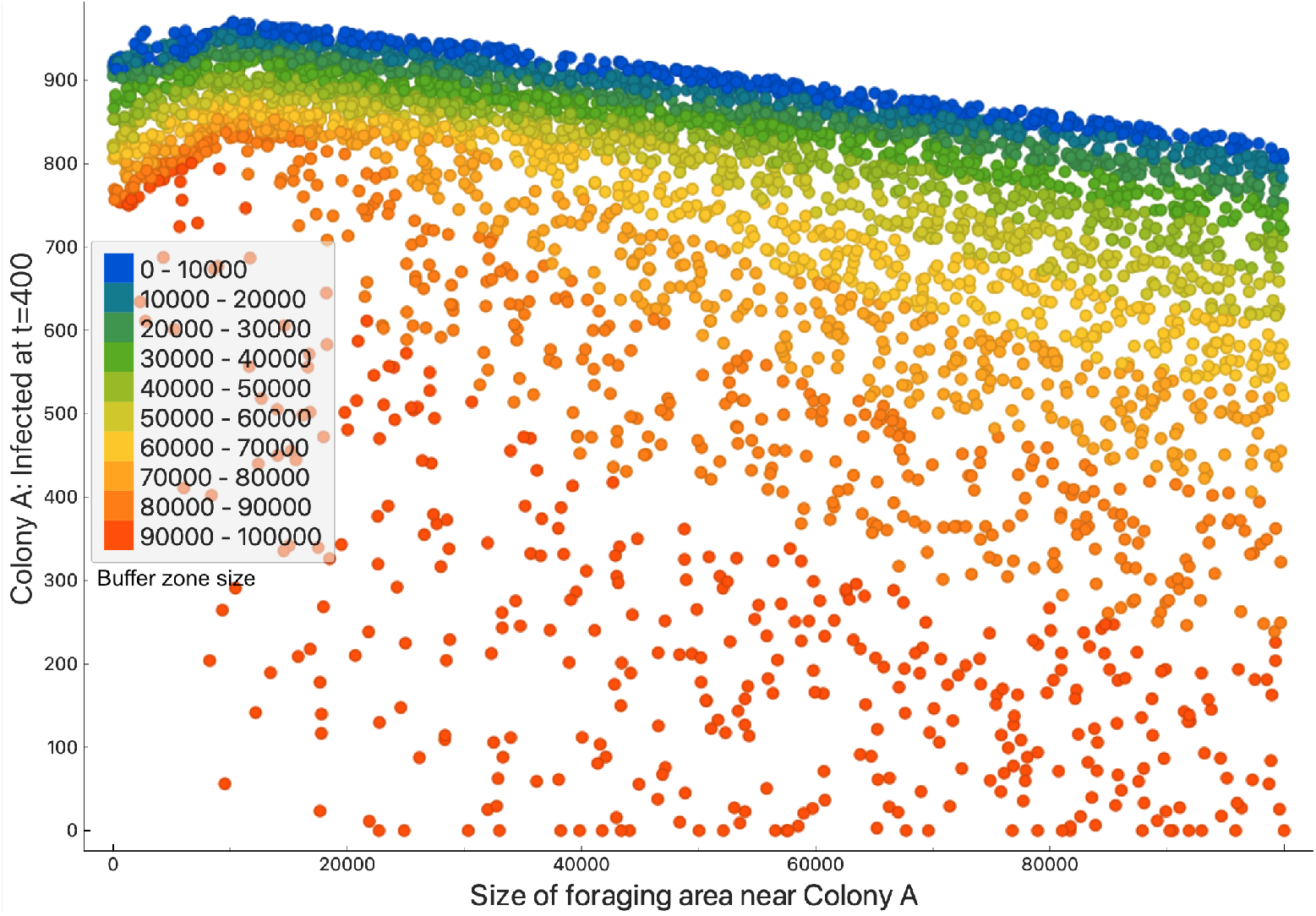
*F*_*A*_ x *I*_*A*_ x *F*_*B*_. Increasing the number of flowers in *F*_*A*_ (size of the foraging area near Colony A) may lead to delayed transmission, particularly if coupled with increased bee pasture in the buffer zone (buffer zone size). The points are generated from 4,000 simulation runs.

**Fig. 8.**
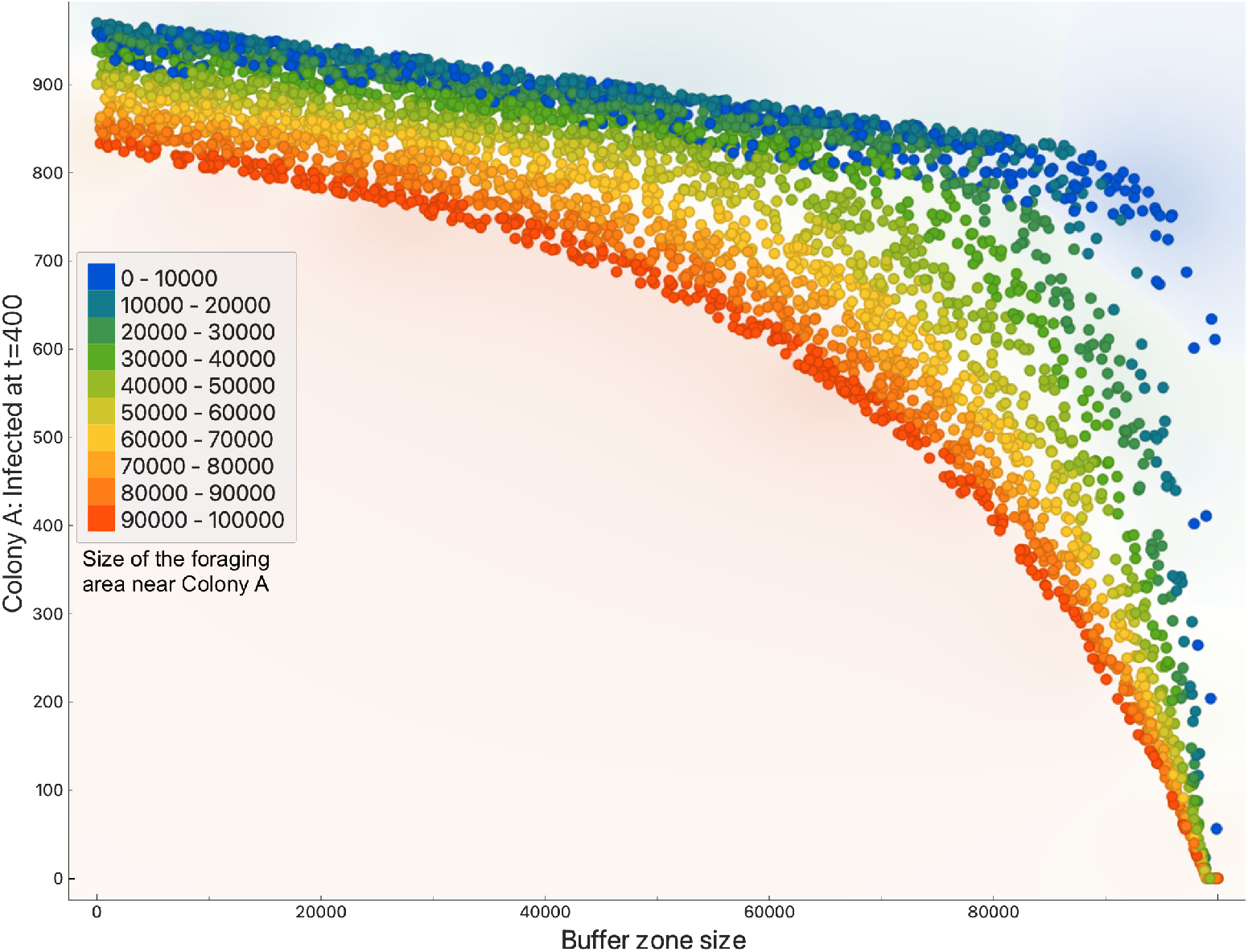
*F*_*B*_ x *I*_*A*_ x *F*_*A*_. The figure vividly illustrates the beneficial impact of increasing the bee pasture in the buffer zone. Bee pasture in *F*_*B*_ can indeed aid in delaying flower-mediated disease transmission, particularly when Colony A has a sufficient food source in *F*_*A*_. The points are generated from 4,000 simulation runs.

## 4 Discussion

As demonstrated in the Results section, sufficient bee pasture in the buffer zone can indeed slow down flower-mediated disease transmission, particularly limiting flower sharing between bees from Colony A and C. This effect is more pronounced when Colony A itself has an adequate food source in its nearby foraging area. Here are some insights and recommendations for beekeepers or anyone aiming to enhance ecological resilience against flower-mediated disease transmission:

1. Ensure there are enough food sources in the nearby area (*F*_*A*_) where Colony A can forage effectively.
2. Establish bee pasture in an outlying area distinct from *F*_*A*_ to serve as the buffer zone, thereby minimizing flower sharing between colonies.
3. Consider additional factors not accounted for in our model, such as seasonality, complex foraging behavior, floral preferences, trait matching, presence of water sources, and beekeeping practices, in future studies to further refine our understanding of disease transmission dynamics.

Relying solely on a buffer zone for flower-mediated disease transmission control is not a foolproof solution, and it does not guarantee complete eradication of infection. It is crucial to complement this strategy with other sound beekeeping practices. A holistic approach that includes regular hive inspections, disease monitoring, and proper hive management practices is essential for maintaining healthy and productive bee colonies [17]. Additionally, promoting biodiversity and providing diverse forage sources can enhance overall colony resilience [41, 42, 49–52]. Nevertheless, the delay in disease transmission facilitated by the buffer zone can provide opportunity and valuable time for beekeepers to plan, prepare, and implement necessary measures to manage bee colonies effectively. This additional time window allows for proactive steps such as relocating hives to safer areas, administering probiotics to strengthen bee health, implementing targeted treatments, or adjusting hive management practices to mitigate the impact of disease outbreaks.

The findings presented in this paper represent one of the pioneering investigations into the role of bee pasture buffer zones in mitigating disease outbreaks. Our results are supported by prior research highlighting the importance of modularity in pollinator-plant interactions. Previous studies have suggested that increased modularity can serve as a protective mechanism, shielding pollinators from flower-mediated pathogen spillover. By compartmentalizing disease outbreaks within distinct modules, the spread of pathogens to other pollinator communities can be inhibited [53]. Furthermore, research indicates that reducing pollinator-plant network connectance, alongside enhancing plant species richness and flower density, may offer benefits to pollinators by effectively partitioning their foraging niches. This partitioning helps limit floral sharing and subsequently reduces the potential for disease transmission. This finding contrasts with the objectives of some pollinator network restoration initiatives, which advocate for increased niche overlap and higher connectance between pollinators and plants to enhance network robustness [54], especially in the face of environmental challenges such as climate change and habitat destruction. Therefore, achieving a balance between these contrasting perspectives is crucial in designing effective foraging areas.

Our proposal to provide ample food sources in nearby foraging areas, coupled with the establishment of a bee pasture buffer zone (such as wildflower strips in outlying areas), offers a solution that reconciles these divergent views. This approach optimizes the benefits of increased modularity and reduced network connectance while minimizing potential drawbacks, ultimately contributing to the resilience and sustainability of pollinator populations.

Managed colonies can also pose a risk of pathogen spillover to wild bee populations [55, 56]. The results presented in this paper extend beyond the protection of managed colonies, such as Colony A, to encompass the conservation of wild bee populations. Consider a scenario where Colony A serves as the source of disease, and wild bees outside the foraging area near Colony A (*F*_*A*_) are at risk of pathogen spillover (Figure 9). In this context, our findings remain applicable: ensuring an adequate food supply near Colony A (*F*_*A*_) restricts the flight of Colony A foragers beyond this area. Moreover, if sufficient food sources are available in the bee pasture buffer zone, wild bees will predominantly visit this area and avoid interaction with Colony A bees in *F*_*A*_. Thus, the establishment of a bee pasture buffer zone emerges as a viable strategy for safeguarding wild bee populations against pathogen spillover, thereby contributing to the conservation of pollinator diversity.

**Fig. 9.**
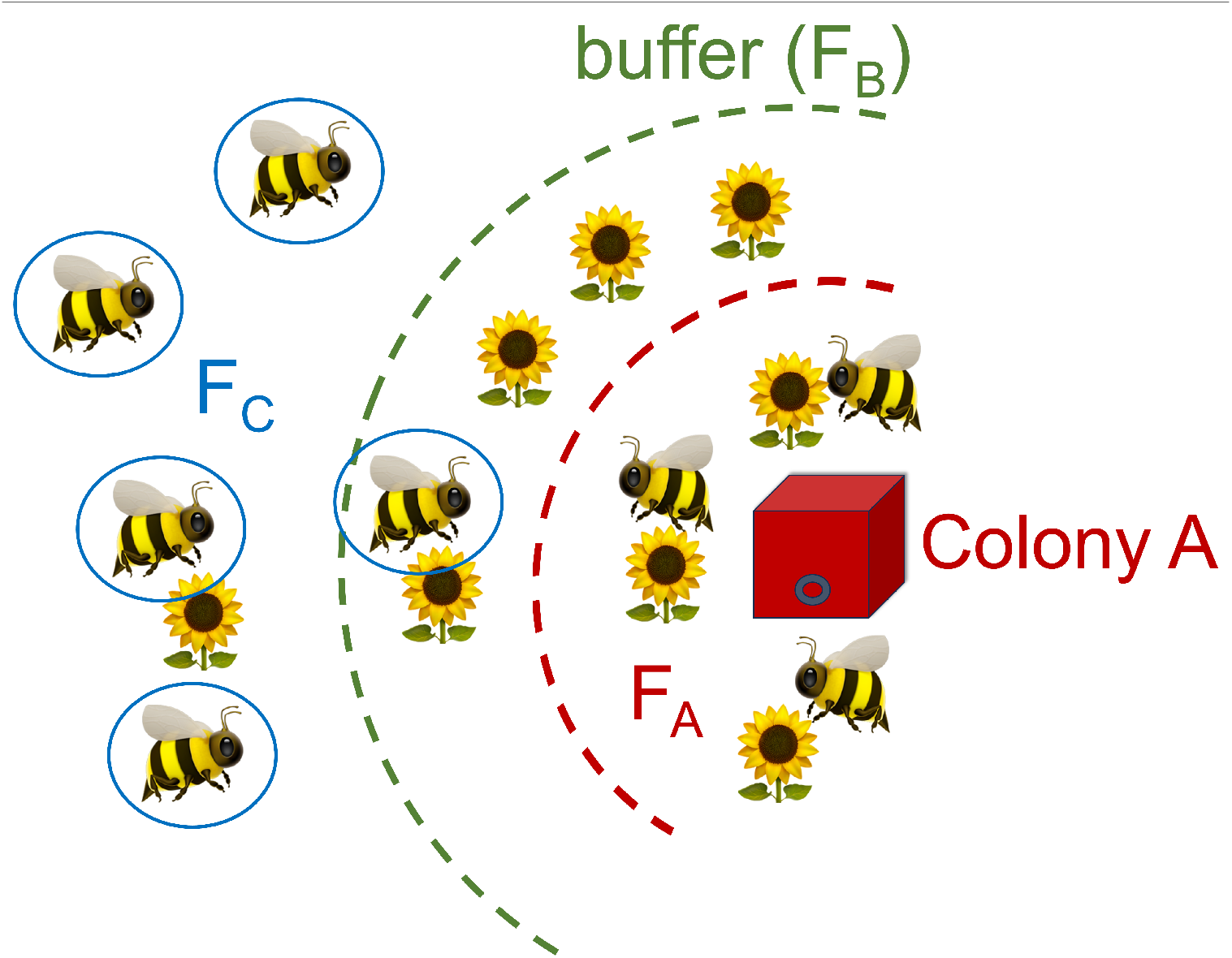
A scenario where managed Colony A serves as the source of disease, and wild bees (encircled in blue) outside the foraging area near Colony A (*F*_*A*_) are at risk of pathogen spillover.

## Acknowledgements

JFR acknowledges support from the Associate Scheme of the Abdus Salam International Centre for Theoretical Physics, Trieste, Italy, and extends gratitude to the UPLB Bee Program for their valuable contributions.

## Supplementary Information

**Table 1.** Model Variables.

| Variables | Definition |
| --- | --- |
| $S_A$ | Population size of Colony A susceptible bees |
| $I_A$ | Population size of Colony A infected bees |
| $I_C$ | Population size of infected Colony C |
| $F_{A,S}$ | Number of flowers in $F_A$ without infectious agents |
| $F_{A,I}$ | Number of flowers in $F_A$ with infectious agents |
| $F_{B,S}$ | Number of flowers in $F_B$ without infectious agents |
| $F_{B,I}$ | Number of flowers in $F_B$ with infectious agents |
| $F_{C,S}$ | Number of flowers in $F_C$ without infectious agents |
| $F_{C,I}$ | Number of flowers in $F_C$ with infectious agents |
| $P_i$ | Function representing new flowers in $F_i$; $i = A, B, C$ |
| $t$ | Time (e.g., minutes) |

**Table 2.** Parameters.

| Parameters | Definition | Values |
| --- | --- | --- |
| $\beta$ | Infectious agent deposit rate in flowers | $[0, 1]$ |
| $\mu$ | Infectious agent removal rate in flowers | $[0, 1]$ |
| $\epsilon$ | Removal rate of flowers | $[0, 1]$ |
| $\alpha$ | Average number of flowers visited by each bee per foraging period | $(0, \infty)$ |
| $\omega_C$ | Increase in $\beta$ when food source is not enough | $[1, \infty)$ |
| $\hat{\beta}$ | Colony A force of infection | $[0, 1]$ |
| $\gamma_i$ | Colony A $F_i$ prioritization coefficient; $i = A, B, C$ | $[0, 1]$ |
| $\omega_A$ | Increase in $\hat{\beta}$ when food source is not enough | $[1, \infty)$ |

### Model Equations

In the mathematical model, there are two important parts. Firstly, we have the equations that represent the transition of flowers from a state without infectious agents to a state with infectious agents. This segment of the model captures the process by which flowers become contaminated as a result of interaction with infected bees from Colony C. Secondly, we have the equations that represent the transition of susceptible bees within Colony A to the infected state. This aspect of the model outlines how susceptible bees become infected through their interaction with flowers harboring infectious agents.

Here are the equations representing the transition of flowers from a state without infectious agents to a state with infectious agents.

*Infectious agent transmission to flowers in foraging area F*_*C*_:

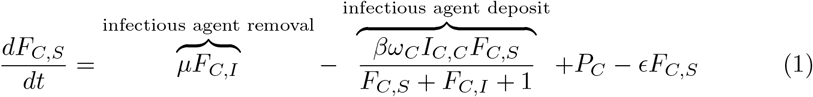

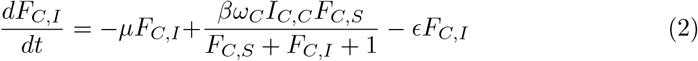

*Infectious agent transmission to flowers in foraging area F*_*B*_:

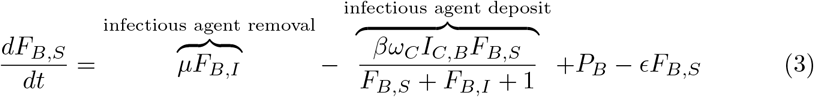

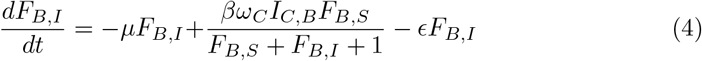

*Infectious agent transmission to flowers in foraging area F*_*A*_:

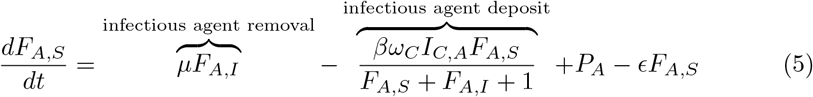

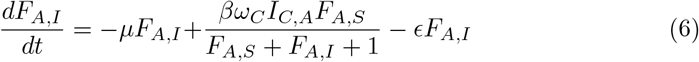

*Number of foragers from Colony C in F*_*C*_, *F*_*B*_ *and F*_*A*_, *respectively, considering floral preference due to distance*:

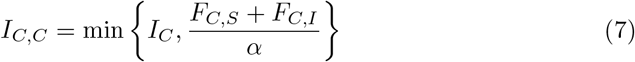

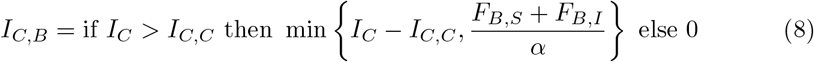

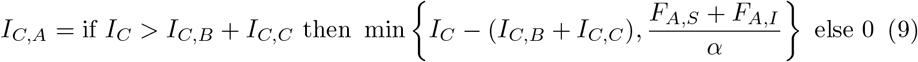

In the above equations, ‘infectious agent removal’ refers to the scenario where infectious agents present in flowers (counted in *F*_*i,I*_, *i* = *A, B, C*) are removed, for example, due to factors such as rain. As a result, these flowers are transitioned back to the state of being without infectious agents and are counted once again in *F*_*i,S*_.

The term ‘infectious agent deposit’ refers to the transition of flowers from a state without infectious agents to a state with infectious agents. This transition occurs due to the visitation of bees from Colony C, represented by *I*_*C,C*_, *I*_*C,B*_ and *I*_*C,A*_. The dilution effect is represented by the denominator.

The parameters *P*_*i*_ and *ϵ* represent the emergence and removal of flowers, respectively. If the simulation time period covers 400 minutes, representing, for example, a 6-hour morning foraging period, the effects of the parameters *P*_*i*_ and *ϵ* could be negligible. In the absence of special events (e.g., transfer of new plants or destruction of flowers by humans or animals), it may be reasonable to set these parameters to zero, as their impact on the model outcomes would be minimal within the given time frame.

The value of *I*_*C,C*_ represents the number of bees from Colony C that can be adequately accommodated by foraging area *F*_*C*_. Each forager has the capacity to visit *α* number of flowers during a foraging period. Similarly, the value of *I*_*C,B*_ denotes the number of bees from Colony C that can be sufficiently accommodated by foraging area *F*_*B*_, excluding those bees already accommodated by *F*_*C*_. Likewise, the value of *I*_*C,A*_ indicates the number of bees from Colony C that can be adequately accommodated by foraging area *F*_*A*_, excluding those bees already accommodated by *F*_*B*_ and *F*_*C*_.

Now, here are the equations representing the transition of susceptible bees within Colony A to the infected state.

*Transmission to Colony A*:

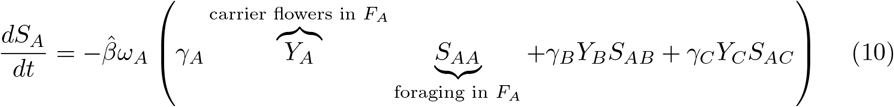

In our results, we monitor the change in the number of infected bees in Colony A (*I*_*A*_) over time, represented by the equation:

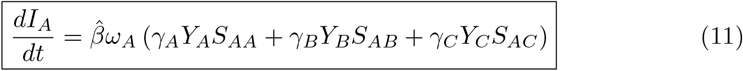

This equation captures the rate of change of infected bees in Colony A (*I*_*A*_) with respect to time (*t*), which depends on various factors including the transmission rate from carrier flower to bee 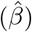, and the foraging dynamics of Colony A with respect to *F*_*A*_, *F*_*B*_, and *F*_*C*_. In particular, the foraging dynamics are influenced by the priorities assigned to different foraging areas, which are based on the distance of Colony A to each foraging area. For instance, Colony A may prioritize foraging in *F*_*A*_ with high priority (e.g., *γ*_*A*_ = 1), followed by *F*_*B*_ with second priority (e.g., *γ*_*B*_ = 0.75), and *F*_*C*_ with least priority (e.g., *γ*_*C*_ = 0.5).

Our objective is to investigate how the number of flowers in the foraging area near Colony A and in the buffer zone influence the number of infected bees in Colony A (*I*_*A*_) after a specified duration, such as *t* = 400 minutes (a 6-hour morning foraging period). This analysis allows us to understand the impact of flower availability in these areas on the dynamics of disease transmission to Colony A.

*Number of flowers with infectious agents in F*_*A*_, *F*_*B*_ *and F*_*C*_ *(carriers), respectively* :

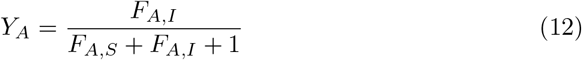

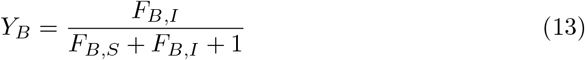

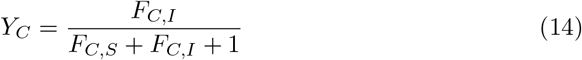

The denominator in these equations (*Y*_*A*_, *Y*_*B*_ and *Y*_*C*_) represent the dilution effect. *Number of susceptible foragers in F*_*A*_, *F*_*B*_ *and F*_*C*_, *respectively* :

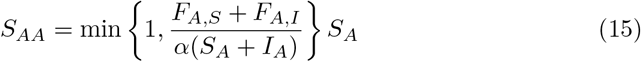

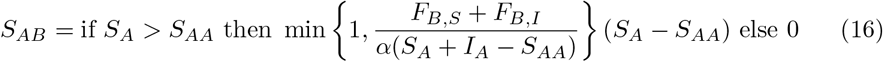

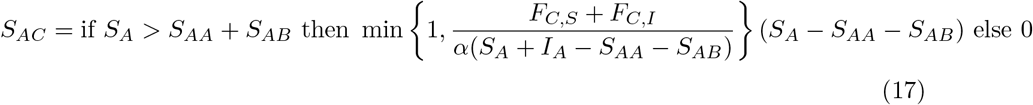

The value of *S*_*AA*_ represents the estimated number of susceptible bees from Colony A that can be adequately accommodated by foraging area *F*_*A*_. Similarly, the value of *S*_*AB*_ denotes the estimated number of susceptible bees from Colony A that can be sufficiently accommodated by foraging area *F*_*B*_, excluding those bees already accommodated by *F*_*A*_. Likewise, the value of *S*_*AC*_ indicates the estimated number of susceptible bees from Colony A that can be adequately accommodated by foraging area *F*_*C*_, excluding those bees already accommodated by *F*_*A*_ and *F*_*B*_.

In addition, there are cases where the flowers in *F*_*A*_, *F*_*B*_ and *F*_*C*_ are insufficient to supply food (e.g., nectar or pollen) for Colony A or Colony C. Here are the equations that represent the increased likelihood of disease transmission resulting from increased flower sharing due to scarcity of floral resources.

*Adjustment in β and* 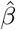, *respectively* :

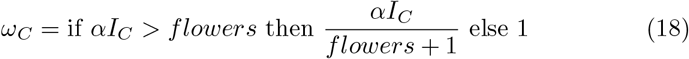

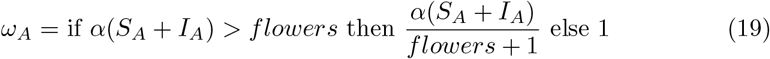

where *flowers* = *F*_*A,S*_ + *F*_*A,I*_ + *F*_*B,S*_ + *F*_*B,I*_ + *F*_*C,S*_ + *F*_*C,I*_.

### Parameter Values

Here are the parameter values used in the simulations presented in the figures within the Results Section. These parameter values were selected to clearly illustrate our point in the paper while maintaining generality. Readers can replicate and validate the generality of our results by numerically solving the model with different combinations of parameter values.

**Table 3.** Parameter values in Figures.

| Parameter | Figure 3 | Figures 4-6 | Figures 7 & 8 |
| --- | --- | --- | --- |
| $S_A(0)$ | 1000 | 1000 | 1000 |
| $I_A(0)$ | 0 | 0 | 0 |
| $I_C(0)$ | Normal(10000,10000/3) | Uniform(100,10000)* | 10000 |
| $F_{A,S}(0)$ | Normal(100,100/3) | Uniform(1000,100000)* | Uniform(0,10000) |
| $F_{A,I}(0)$ | 0 | 0 | 0 |
| $F_{B,S}(0)$ | x-axis in the Figure | Uniform(1000,100000)* | Uniform(0,10000) |
| $F_{B,I}(0)$ | 0 | 0 | 0 |
| $F_{C,S}(0)$ | Normal(10000,10000/3) | Uniform(1000,100000) | 1000 |
| $F_{C,I}(0)$ | 0 | 0 | 0 |
| $P_i$ | 0 | 0 | 0 |
| $\beta$ | Normal(0.1,0.1/3) | Uniform(0.01,0.1) | 0.1 |
| $\mu$ | Normal(0.01,0.01/3) | Uniform(0.01,0.2) | 0.01 |
| $\epsilon$ | 0 | 0 | 0 |
| $\alpha$ | Normal(10,10/3) | Uniform(0.5,10) | 10 |
| $\hat{\beta}$ | Normal(0.01,0.01/3) | Uniform(0.01,0.1) | 0.01 |
| $\gamma_A$ | 1 | 1 | 1 |
| $\gamma_B$ | Uniform(0.5,1) | Uniform(0.5,1) | 0.75 |
| $\gamma_C$ | Uniform(0.1,1) | Uniform(0.1,1) | 0.5 |

\*except when the values are in the x-axis of the Figure
Note: Each selected value from Normal(mean,standard deviation) or Uniform(start,end) will be used consistently throughout one simulation run.

